# VIPERA: Viral Intra-Patient Evolution Reporting and Analysis

**DOI:** 10.1101/2023.10.24.561010

**Authors:** Miguel Álvarez-Herrera, Jordi Sevilla, Paula Ruiz-Rodriguez, Andrea Vergara, Jordi Vila, Pablo Cano-Jiménez, Fernando González-Candelas, Iñaki Comas, Mireia Coscollá

**Affiliations:** Institute for Integrative Systems Biology (I2SysBio, University of Valencia - CSIC), FISABIO Joint Research Unit “Infection and Public Health”, Paterna, Spain; Department of Clinical Microbiology, CDB, Hospital Clínic of Barcelona; University of Barcelona; ISGlobal, Barcelona, Spain; CIBER of Infectious Diseases (CIBERINFEC), Madrid, Spain; Institute of Biomedicine of Valencia (IBV-CSIC), Valencia, Spain; CIBER of Epidemiology and Public Health (CIBERESP), Madrid, Spain

**Keywords:** SARS-CoV-2, within-host evolution, serially-sampled infection, intra-patient diversity, Snakemake workflow, bioinformatics

## Abstract

Viral mutations within patients nurture the adaptive potential of SARS-CoV-2 during chronic infections, which are a potential source of variants of concern. However, there is no integrated framework for the evolutionary analysis of intra-patient SARS-CoV-2 serial samples. Herein we describe VIPERA (Viral Intra-Patient Evolution Reporting and Analysis), a new software that integrates the evaluation of the intra-patient ancestry of SARS-CoV-2 sequences with the analysis of evolutionary trajectories of serial sequences from the same viral infection. We have validated it using positive and negative control datasets and have successfully applied it to a new case, thus enabling an easy and automatic analysis of intra-patient SARS-CoV-2 sequences.

## Background

During the severe acute respiratory syndrome coronavirus 2 (SARS-CoV-2) pandemic, almost 7 million deaths have been reported by the World Health Organization (WHO) [1] due to COVID-19. The pandemic has been driven by SARS-CoV-2 variants of concern (VOC), which are variants with an increased pathogenicity [2]. This VOCs have appeared several times in the COVID-19 pandemic, and it has been observed that the clades containing the VOCs are preceded by a steam branch that shows, on average, a 4-fold increase in the substitution rate [3], which was usually around 10^-3^ substitutions per site and year in 2020 [4,5].

Different hypotheses —such as undetected acute infections [6] or secondary hosts— have been proposed to explain the increase in the substitution rate and thus, the appearance of VOCs. Nowadays, several pieces of evidence support the hypothesis that VOCs originated in chronic infections. First, the immune system of immunocompromised patients can fail to clear acute SARS-CoV-2 infections leading to long term infections [7]. The high number of viral mutations from long term infections, most of them in the spike protein coding region [8], would suggest an increased evolutionary rate, as observed in branches that give rise to VOCs clades [9]. Second, defining mutations of several VOCs have been detected in sequences from chronic infections [10]. Following these findings, there has been an effort to study SARS-CoV-2 chronic infections, trying to enhance the surveillance of VOCs, but also to better understand the mechanisms behind their emergence [8,11–13]. While there are pipelines that integrate reproducible workflows to analyze genomic diversity between patients [14,15], there is a lack of easily deployable, accessible, and integrated workflows for analyzing and reporting the evolutionary trajectories of SARS-CoV-2 chronic infections. Current pipelines for processing serially-sampled sequencing data that take into account the particularities of intra-host samples are restricted to certain analyses, such as detecting mixed viral populations, or identifying chronic infections but using only consensus sequences [12,16–19]. For this reason, carrying out this type of studies through public databases is a difficult task especially without further clinical information.

Here, we present VIPERA (Viral Intra Patient Evolution Reporting and Assessment), a user-friendly workflow to easily identify and study within-host evolution in SARS-CoV-2 serially-sampled infections. Our tool provides an aggregate of population genomics and phylogenetic analyses that allows researchers to determine if a collection of SARS-CoV-2 samples originates from a serially-sampled viral infection. Furthermore, VIPERA provides insights into intra-host evolution, tracking variant trajectories and selective pressure over time.

## Results

### A comprehensive report of a serially-sampled SARS-CoV-2 infection

VIPERA offers an integrated framework for detecting and studying serially sampled SARS-CoV-2 infections. The necessary data inputs are the read mappings (in BAM format) and the consensus genomes (in FASTA format) for each sequence of the target dataset, as well as the associated sample metadata. The main output from VIPERA is a report file in HTML format summarizing all the analyses in three main sections: “1. Summary of the target dataset”, “2. Evidence for single, serially-sampled infection”, and “3. Evolutionary trajectory of the serially-sampled SARS-CoV-2 infection”. In addition, the intermediate files which are instrumental in the creation of the final report —such as the lineage demixing summary, the maximum-likelihood phylogeny of the target dataset within its spatiotemporal context, the pairwise weighted-distance matrix for the target dataset, or the variant calling results with the dataset ancestor as reference— are also made available to the user (see Additional file 1: Table S1 for a full list). This offers a great degree of flexibility and control over the data, allowing for further in-depth analysis if required. The three sections of the report are described hereafter.

#### 1. Summary of the target sample dataset

First, the report displays a summary of the target sample dataset that includes the date and location of sampling. This summary also reports the lineage assignment and a time-sorted index of each sample that is used to identify the samples in the downstream analyses.

#### 2. Evidence for single, serially-sampled infection

The first aim of VIPERA is to streamline the process of confirming that samples originate from a single, serially-sampled infection collected from the same patient at different time points —as opposed to multiple successive infections, co-infections, or instances of sample contamination. For this, the following analyses are conducted.

##### 2.1. Lineage admixture

A lineage composition profile of each sample based on read mappings is reported to detect if different viral lineages are present in the sample (e.g. in co-infections or contaminations).

##### 2.2. Phylogeny and temporal signal

A maximum-likelihood tree including target and context samples is displayed in the VIPERA output. A group of SARS-CoV-2 sequences originating from a serially-sampled infection must be monophyletic. The phylogeny enables users to assess whether the target samples are monophyletic based on ultrafast bootstrap (UFBoot) and the Shimodaira–Hasegawa-like approximate likelihood ratio test (SH-aLRT) support values.

Additionally, the temporal signal is also evaluated for the studied samples. When previous evidence supports the hypothesis, a robust temporal signal further validates that the target dataset was serially sampled from a single infection.

##### 2.3. Nucleotide diversity comparison

The nucleotide diversity (π) for the target samples is compared with the distribution of π obtained for random subgroups extracted from a patient-independent context dataset. If the target dataset has a significantly lower π than the distribution of π values for sequences from different patients, then we can assume that they come from the same viral infection. The report includes the estimated significance of π being lower in the target samples.

#### 3. Evolutionary trajectory of the serially-sampled SARS-CoV-2 infection

The next step is to characterize within-host evolution. To this end, VIPERA reports a set of analyses focused on describing the intra-host evolutionary trajectory of the target samples.

##### 3.1. Number of polymorphic sites

To investigate the within-host viral diversity we use the number of polymorphic sites (minor allele frequency > 0.05) as a measure of diversity. The report displays the number of polymorphic sites of each sample and the correlation of this parameter with time, which allows for the observation of fluctuations in diversity throughout the course of the infection.

##### 3.2 Description for within-host nucleotide variants

The report includes a summary of within-host nucleotide variants with respect to its predicted ancestral sequence. The summary includes a genome-wide depiction of the proportion of sites in which we find a polymorphism. This allows for the identification of mutation hotspots. The summary also depicts each individual mutation throughout the genome for each sample. Mutations are represented according to their classification in single-nucleotide variants (SNVs) or insertions and deletions (indels) and colored depending on whether they are synonymous or non-synonymous SNVs, in-frame or frameshift indels, or intergenic nucleotide changes. Due to the relevance of the spike protein for SARS-CoV-2 adaptation, a zoom-in of the summary is also generated for the S gene.

##### 3.3. Time dependency for the within-host mutations

Allele frequencies at each polymorphic site are tested for correlation with time. In the report, the correlation coefficient and the adjusted significance of the correlation is included first. Then, significantly positively correlated allele frequencies — assumed to be affected by selective pressures or hitchhiking— are displayed on a time series of allele frequencies, along the viral genome. All sites with more than one alternative allele are also displayed.

##### 3.4. Correlation between alternative alleles

To evaluate if there are interactions between mutations, the report includes an interactive heatmap of pairwise allele frequency correlation coefficients, which includes the relationships between alleles. The interactive heatmap enables the user to easily obtain correlation values and restrict the region for visualization.

##### 3.5. Non-synonymous to synonymous rate ratio over time

Finally, the report includes a time series of the synonymous mutations per synonymous site (dS) and non-synonymous mutations per non-synonymous site (dN) of each sample with respect to the ancestor sequence.

### Validating the detection of serially-sampled infections

To validate the evidence of serially-sampled infection we tested the pipeline with two control sets of samples. The positive control dataset includes 30 sequences from a chronic infection collected in Yale between February 8, 2021, and March 7, 2022 [11]. All sequences from the positive control were designated as the B.1.517 lineage. Its context dataset (n = 170) was automatically fetched from GISAID, searching for samples assigned to the same lineage, and collected in the same location, from February 1, 2021, to March 12, 2022.

The negative control dataset combines 15 sequences from two different patients (4:1 ratio). Both were collected in Barcelona between March 24, 2020, and November 16, 2020, and designated as lineage B.1 (see Material and Methods). Its context dataset (n = 84) was also automatically fetched from GISAID by searching for the same lineage, and collected in the same location, from March 11, 2020, to November 28, 2020.

### Lineage composition analysis

When samples were decomposed in lineages, two different landscapes appeared in the positive and negative control datasets. All 30 samples from the positive control had a 100% estimated abundance of the B.1.517 lineage (Figure 1A). Conversely, for the negative control, five samples were mostly B.1 or B.1.399, while in the remaining 10 samples, B.1 and B.1 sublineages had an estimated abundance of up to 88% (Figure 1B).

**Figure 1.**
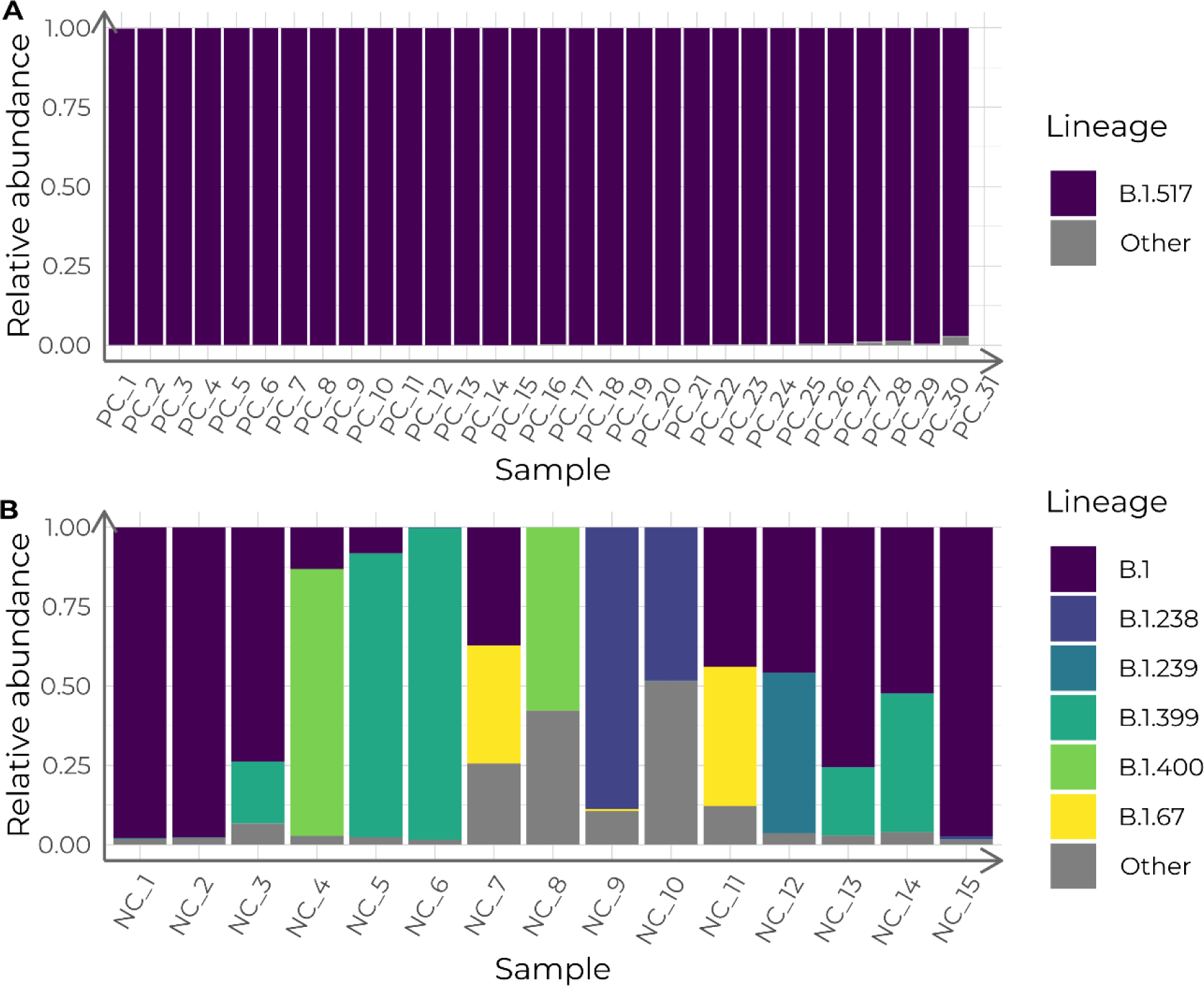
**Lineage admixture of the control datasets, calculated with Freyja**. Columns depict the estimated relative lineage abundance in each sample in the positive control (PC) dataset (A) and in the negative control (NC) dataset (B). Samples in the X-axis are ordered chronologically, from more ancient to newer.

### Monophyly supports the detection of serially-sampled infections

A maximum-likelihood tree was constructed with both the target and the context datasets for the two validation cases. In the positive control, all 30 samples fell into a robust clade together with other eight sequences from the context dataset (UFboot: 97 %; SH-aLRT: 77 %) (Figure 2A). Those eight samples were later confirmed to have been sampled from the same patient (personal communication with Dr. Anne Hahn and Dr. Nathan Grubaugh). Thus, considering the eight additional sequences as part of our study dataset, rather than part of the context, we can conclude that the positive control sequences were monophyletic.

**Figure 2.**
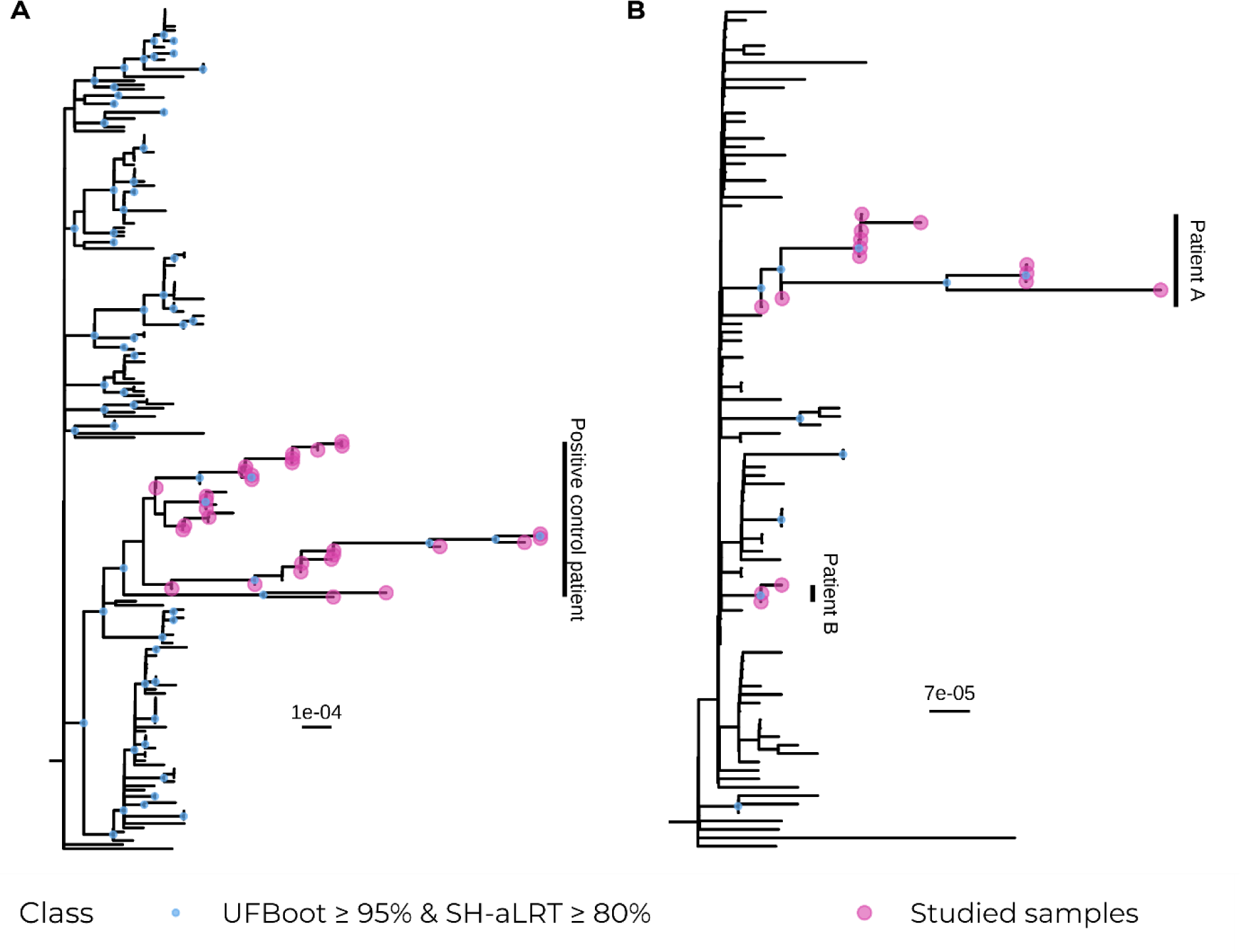
**Maximum-likelihood phylogenies of the control datasets and their context samples with 1000 support replicates**. Positive control dataset. B) Negative control dataset.

As for the negative control, all 15 sequences were paraphyletic and fell into a clade with weak support (UFBoot: 7.0 %; SH-aLRT: 0.00 %) together with another 61 context sequences. However, sequences were divided into two strongly supported monophyletic clades that correspond with the two groups of samples coming from two different patients that we had artificially mixed. One clade contained the 3 sequences from the patient B of the negative control (UFBoot: 96 %; SH-aLRT: 92 %) and the other clade contained the 12 sequences from the patient A of the negative control (UFBoot: 97 %; SH-aLRT: 87 %) (Figure 2B).

Based on the pairwise distance between samples accounting for allele frequencies, neighbor-joining trees were constructed for each control dataset (Figure 3A and Figure 3C). Root-to-tip distances were used to estimate their temporal signal (Figure 3B and Figure 3D). We found a robust temporal signal for the positive control dataset, with an estimated 24.94 substitutions per year, 95% confidence interval (CI) [19.59, 30.28] (R^2^ = 0.76, F(1, 28) = 91.26, p < 0.001; Figure 3B). In light of previous evidence supporting the dataset having been serially sampled from an intra-patient infection, the temporal signal further supported the hypothesis. Additionally, we found a robust temporal signal in the negative control dataset too, with an estimated 30.82 substitutions per year, 95% CI [25.21, 36.43] (R^2^ = 0.92, F(1, 13) = 141.1, p < 0.001; Figure 3D). Since earlier findings did not back up the serial sampling scenario, the temporal signal does not hold any value as evidence for the hypothesis for the negative control dataset.

**Figure 3.**
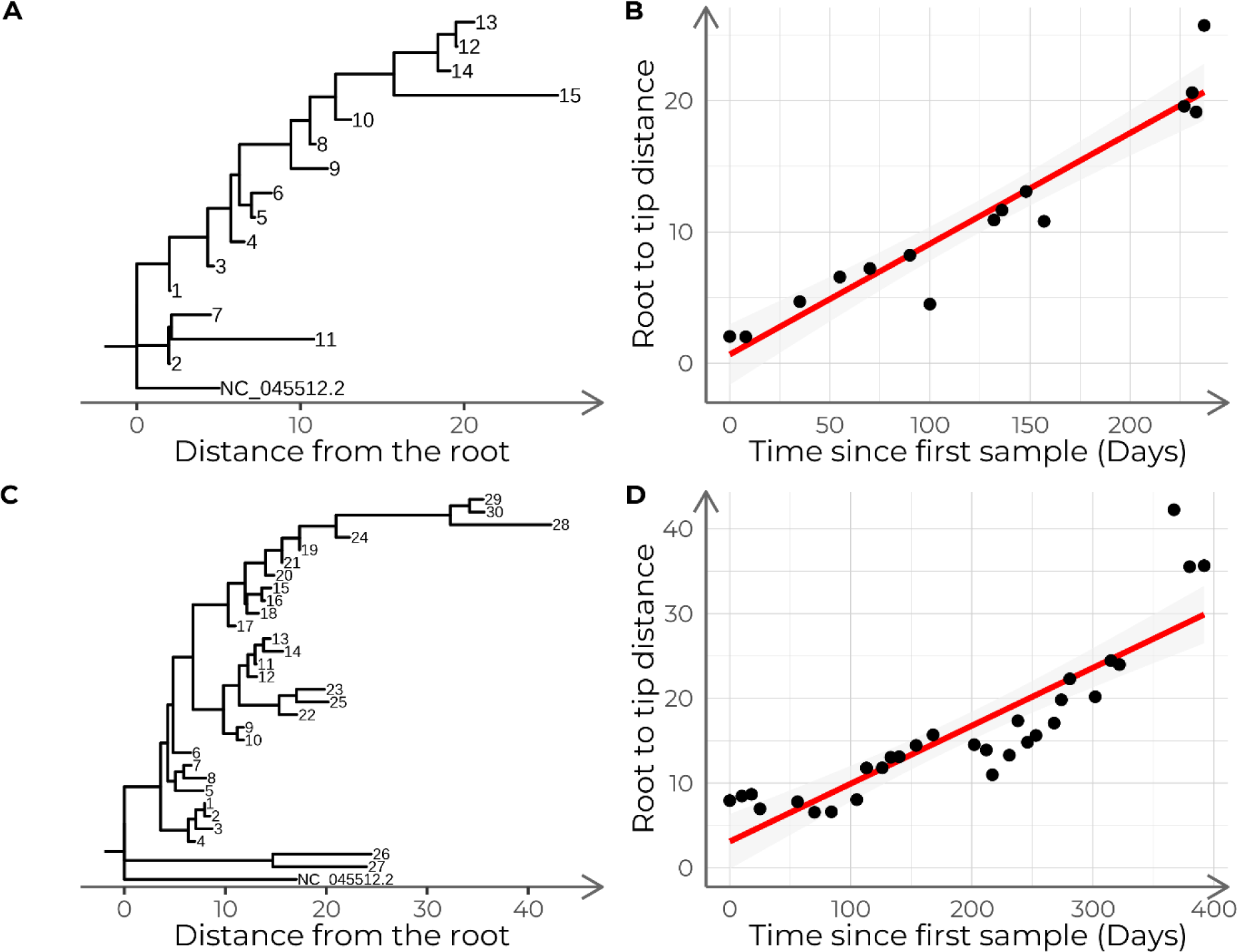
**Neighbor-joining trees of the control datasets and time series of tree root-to-tip distances**. Trees are based on pairwise allele frequency-weighted distances and include the samples that compose the negative control (A) and the positive control (C). The scatterplot shows the relationship between root-to-tip distances and the number of days passed since the first sample for the positive control (B) and the negative control (D). The red lines depict the linear model fit.

### Nucleotide diversity reveals chronic infections

For each validation dataset, we calculated the nucleotide diversity of the studied samples and compared it with the nucleotide diversity of 1000 subsets of samples of the same size as the target dataset, extracted from each corresponding context dataset. The nucleotide diversity of the positive control (π = 1.80·10^-4^) was significantly lower than that of its corresponding context dataset (average = 5.30·10^-4^, SD = 2.87·10^-5^; t-test t = 376.27, p < 0.001; Figure 4A) assuming a normal distribution of the context π values (Shapiro-Wilk test W = 0.997, p = 0.076). Conversely, the negative control dataset did not show a significantly lower nucleotide diversity (π = 1.03·10^-4^) compared to its context dataset π distribution (average = 1.34·10^-4^, SD = 3.55·10^-5^; empirical p = 0.137; Figure 4B) without assuming normality (Shapiro-Wilk test W = 0.98, p < 0.001).

**Figure 4.**
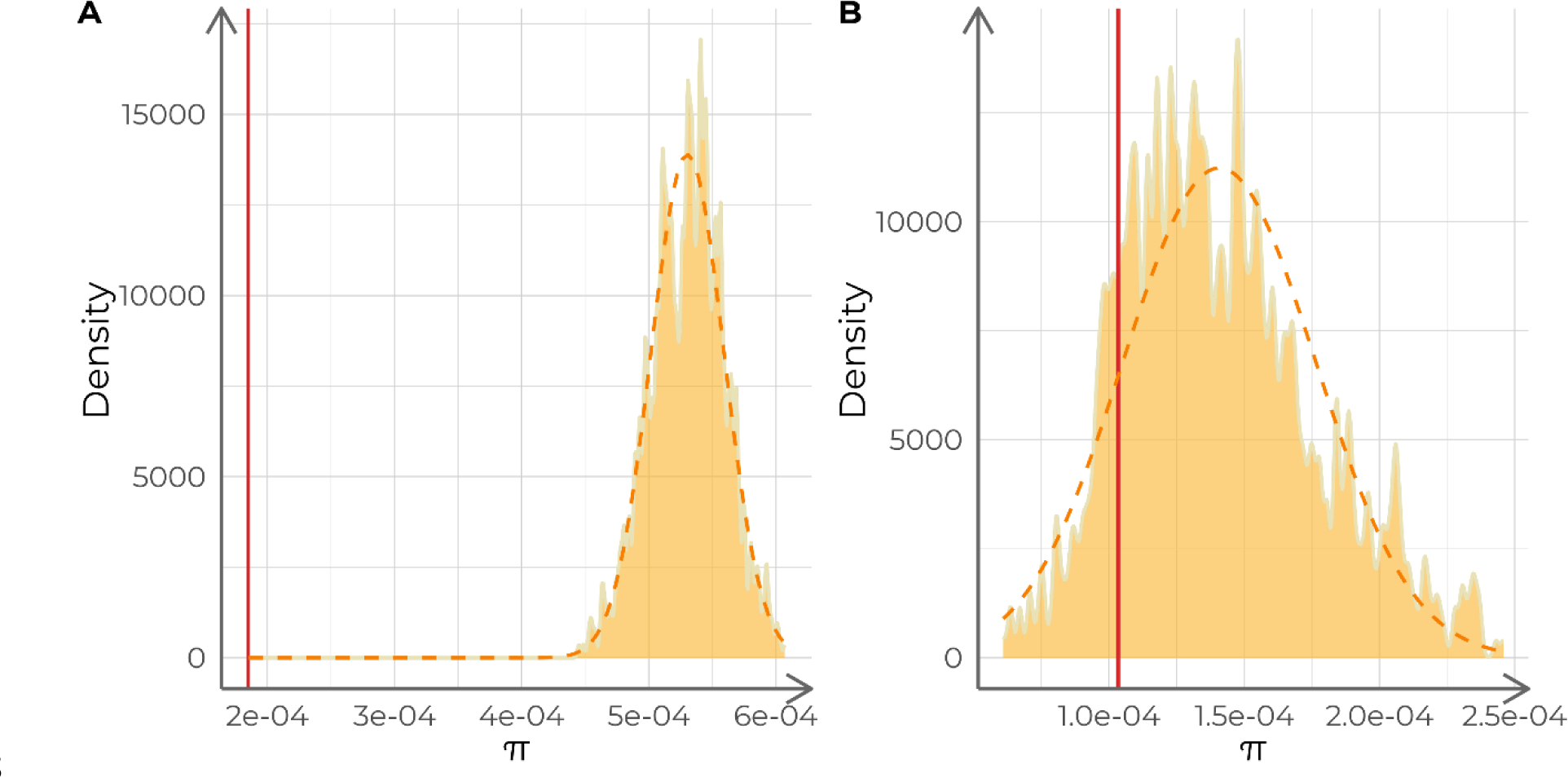
**Analysis of the nucleotide diversity (π) of each control dataset**. The orange dashed lines describe a normal distribution with the same mean and standard deviation as the distribution of π values. The red vertical lines indicate the π value for the studied samples. A) Analysis of the positive control against 1000 replicates (n = 15 each) of its context dataset. Analysis of the negative control against 1000 replicates (n = 30 each) of its context dataset.

Furthermore, we repeated the analysis of the positive control, but considered the eight additional samples of the same patient as a part of the studied samples, instead of the context. Nucleotide diversity was lower compared with the original analysis (π = 1.3·10^-4^). Additionally, it was significantly lower compared to its corresponding context (average = 5.20·10^-4^, SD = 2.45·10^-5^; t-test t = 514.19, p < 0.001).

### Using VIPERA to analyze a novel case

We applied the pipeline to study the within-host evolution in a set of 12 SARS-CoV-2 samples collected from the same patient and designated to lineage B.1. These 12 sequences belong to patient A included in the negative control. Their context dataset was automatically constructed searching for B.1 sequences collected in Barcelona between March 24, 2020, and November 16, 2020, in the GISAID database, and included 85 sequences. Additionally, another custom context dataset was also constructed with 110 samples manually selected from the SEQCOVID Consortium. These were collected in Barcelona from independent patients between March 11, 2020, and November 28, 2020, and classified as B.1. Results using both context datasets were consistent, so we report those with the automatically constructed context dataset because it is the default VIPERA option.

### Evidence for single, serially-sampled infection

#### Weakly defined lineages can lead to false lineage admixtures

We investigated the most probable lineage admixture for all 12 samples. We observed two pairs of samples with an estimated lineage abundance of nearly 100% for lineages B.1 and B.1.399, respectively.

The remaining samples were further classified in B.1 sublineages, with their estimated abundances ranging from 0.07% to 88% (Figure 5A). The low number of mutations between B.1 and B.1 sublineages (1-2 SNPs) might reflect variations during the evolution of the virus over time rather than the mixture of different viruses.

**Figure 5.**
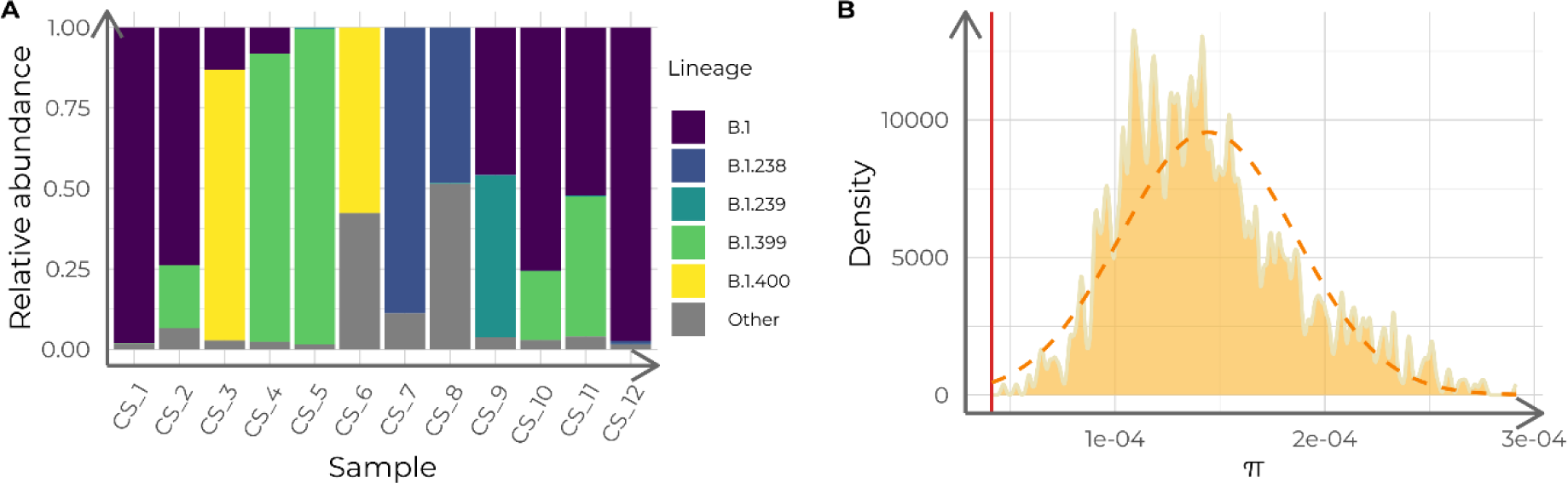
**Lineage admixture and nucleotide diversity (π) analysis of the 12 case study samples**. A) Estimated relative lineage abundance in each of the 12 case study samples, calculated with Freyja. Samples in the X-axis are time-ordered from more ancient to newer. B) Nucleotide diversity (π) distribution for 1000 samples (n = 12) of context sequences for the case study. The orange dashed curve depicts a normal distribution with the same mean and standard deviation as the π value distribution. The red vertical line indicates the π of the case study dataset.

#### All target samples form a monophyletic cluster

The maximum-likelihood phylogeny revealed that the case study dataset formed a monophyletic cluster. The clade that contained all studied samples was supported by a UFBoot score of 97 % and a SH-aLRT score of 92 % (Figure 6A and 6B).

**Figure 6.**
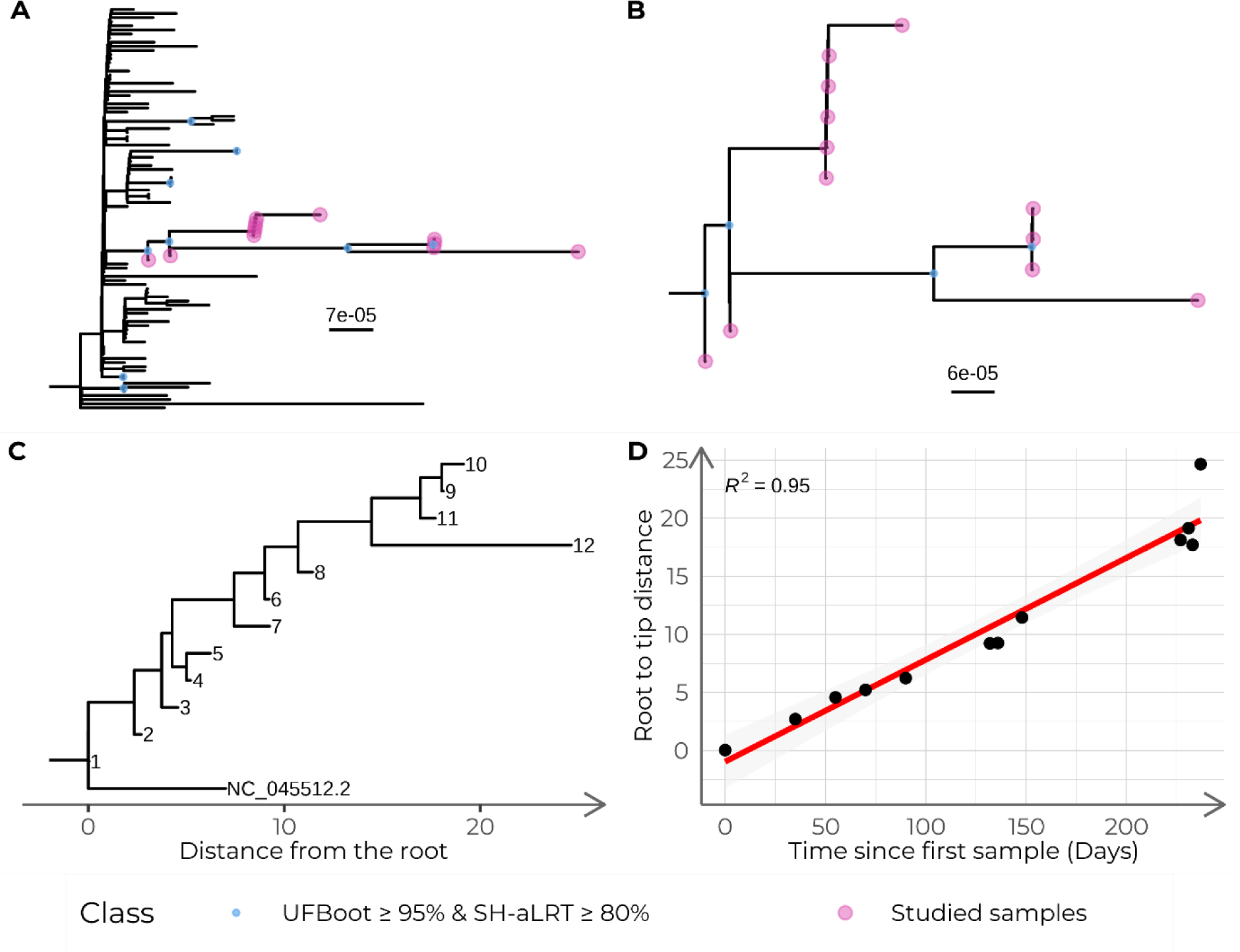
**Phylogenetic analysis of the case study dataset**. A) Maximum-likelihood phylogeny with 1000 supporting replicates for both studied and context samples of the case study. The clade containing all target samples is highlighted in red. B) Zoom of the clade in (A) containing all studied samples. C) Neighbor-joining tree constructed with pairwise weighted distances for the case study samples. D) Temporal signal for the case study using a neighbor-joining tree constructed with pairwise weighted distances. The red line depicts the linear model fit.

Allele frequency-weighted pairwise distances were calculated, and a neighbor-joining tree was constructed (Figure 6C). Time (in days) since the first sample predicted root-to-tip distances (R^2^ = 0.95, F(1, 10) = 174.8, p < 0.001) with an estimated substitution rate of 32.02 substitutions per year, 95% CI [26.62, 37.41] (Figure 6D).

#### Nucleotide diversity is reduced when compared with context samples

The nucleotide diversity (π = 4.11·10^-5^) was lower than that of its corresponding context dataset (average = 1.44·10^-4^, SD = 4.04·10^-5^; empirical p < 0.001; Figure 5B) without assuming a normal distribution of the context π values (Shapiro-Wilk test W = 0.967, p < 0.001). This finding supports the hypothesis of these sequences coming from a serially-sampled, single-virus infection.

To summarize the evidence from section 2 of the report. Firstly, we found that lineage composition analysis supported a homogeneous lineage classification of all serial samples. Secondly, the maximum-likelihood phylogeny showed that the studied samples are monophyletic, thus indicating a proximal common origin. Thirdly, the analysis of nucleotide diversity showed that it was significantly lower in the studied dataset than in the context dataset. Finally, the strong temporal signal observed in the studied samples in addition with the previous evidence led us to conclude the common infectious origin of the serially sampled studied samples. Based on this premise, we proceeded to examine intra-host evolution, which is described in the following part of the report.

### Evolutionary trajectory of the serially-sampled SARS-CoV-2 infection

#### Diversity increases over time

Using the number of polymorphic sites as an estimate of genetic diversity, we observed that diversity was positively correlated with time in days since the first sample (Figure 7A). In fact, time since the initial sampling significantly predicted the number of polymorphic sites (R^2^ = 0.7, F(1, 10) = 22.69, p < 0.001).

**Figure 7.**
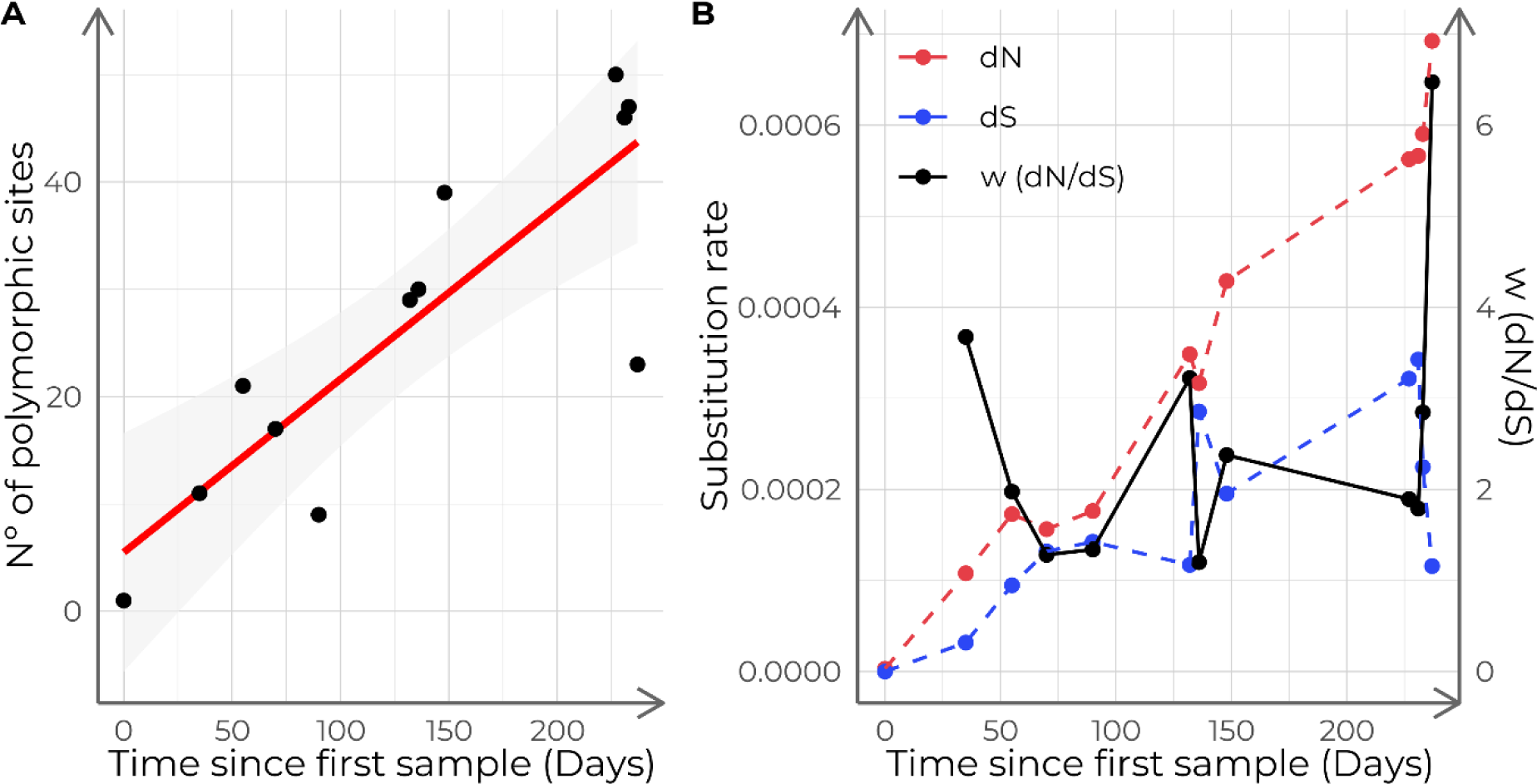
**Diversity analysis of the case study samples**. A) Number of polymorphic sites of the case study samples, depending on collection date. The red line shows the linear model fit. B) Time series of dN and dS and ω (dN/dS). Each point corresponds to a different sample, sorted in chronological order.

#### Nucleotide variants appearing due to within-host evolution

We found 10 indels, six of which led to frameshifts: 2 in the ORF1ab, 2 in the ORF7b, one in the ORF3a and other in the N gene. Also 99 different SNVs were found, 67 of which were non-synonymous (see Additional file 2). Genomic variation was not evenly distributed along the SARS-CoV-2 genome. Some regions such as NSP3 in the ORF1ab, the S gene and the N gene reached peaks of 1% of polymorphic sites (Figure 8).

**Figure 8.**
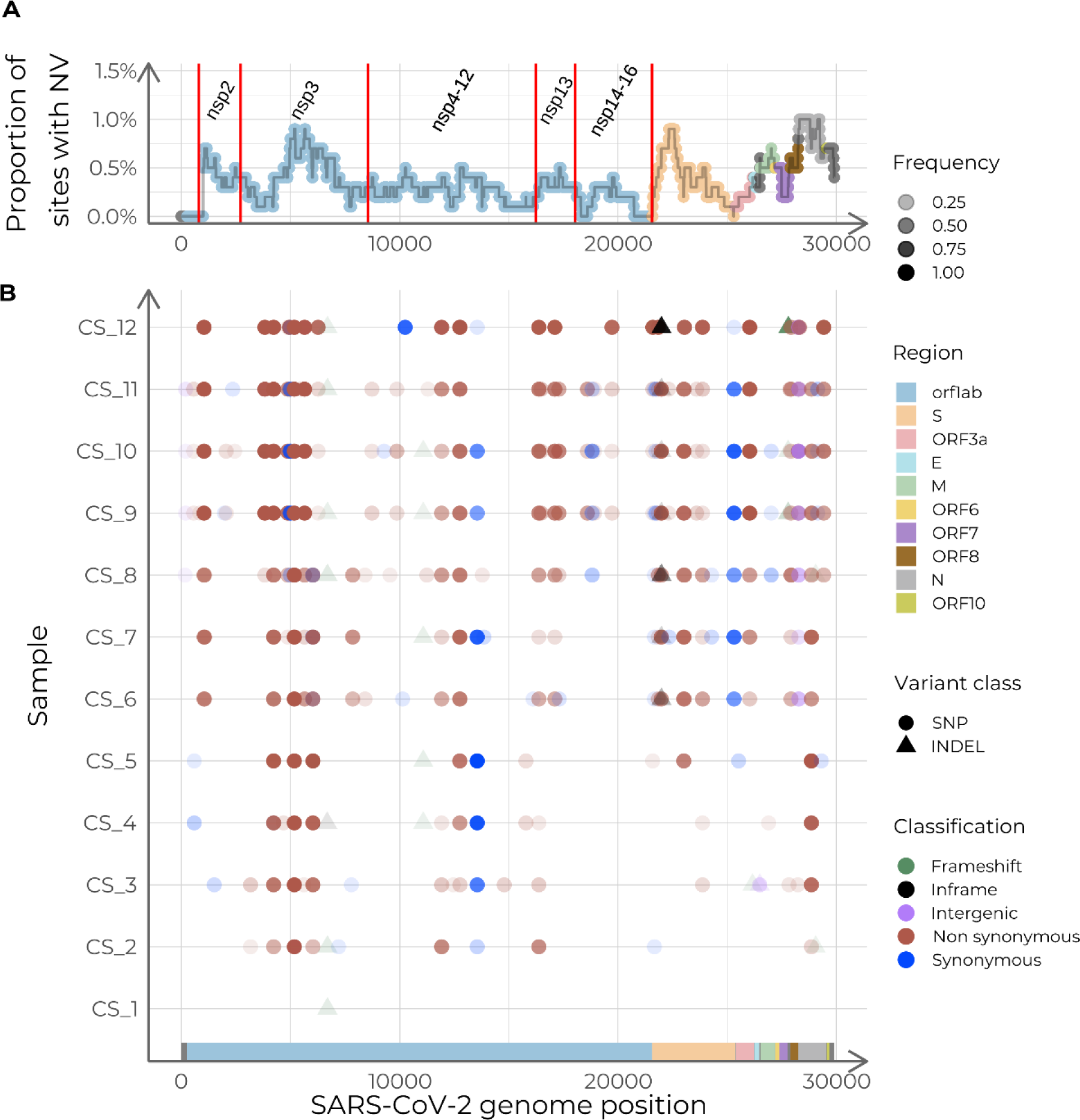
**Summary of the intra-host accumulation of nucleotide variants (NV), using the dataset ancestor as reference**. A) Nucleotide variants per site along the SARS-CoV-2 genome. Relative abundance of NVs is calculated with a sliding window of width 1000 nucleotides and a step of 50. Labels indicate the coding regions of the non-structural proteins (NSP) within ORF1ab. B) Genome variation along the genome for each sample. The Y-axis displays samples in chronological order, with the earliest collection date at the bottom, and the latest, at the top.

The nucleotide variants found were tested for their correlation with time. Eight out of 109 showed a significant correlation with time, being positive for all of them, with Pearson’s coefficients ranging from 0.873 to 0.957 (Figure 9A). We also found two positions with more than one alternative allele (Figure 9). Site 4230 had one allele that was positively correlated with time (ORF1ab:T1322K and ORF1ab:T1322I, both located in the coding region of NSP3). Two deletions in the S gene were detected: S:V143D + ΔY144 and S:V143D + ΔY144/Y145 (Δ21990-21992 and Δ21990-21995 at the genome level, respectively).

**Figure 9.**
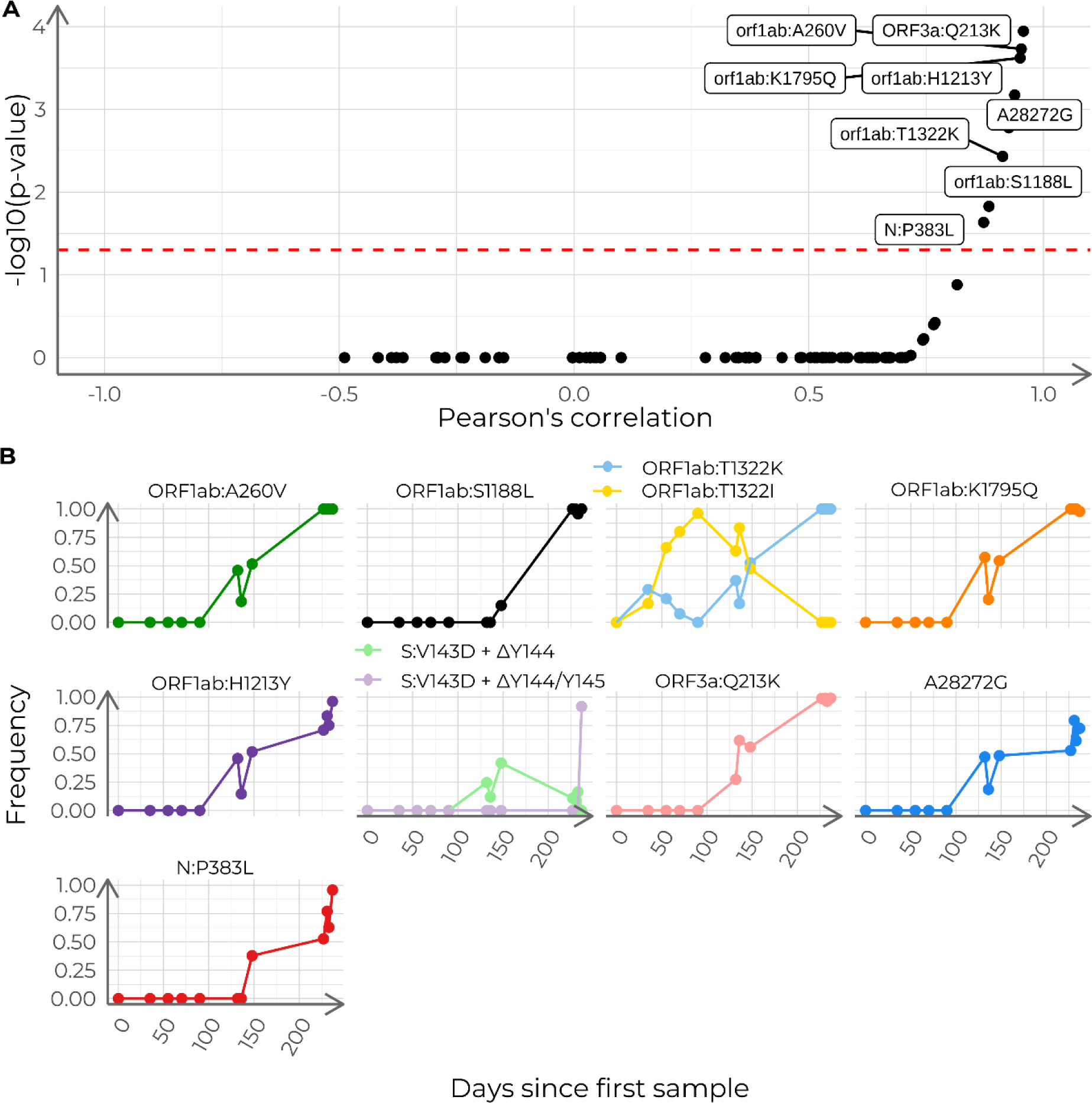
**Analysis of the accumulation of polymorphisms in the case study**. A) Pearson’s correlation coefficients and BH-adjusted p-values for all 110 detected nucleotide variants. Red dashed line indicates adjusted p = 0.05. Labeled dots represent nucleotide variants correlated with time (adjusted p < 0.05). B) Time series of relative allele frequencies. The shown positions include nucleotide variants with a significant correlation with time and sites with more than two possible states. Each subplot depicts the progression of the allele frequencies in time for a given genome position.

Moreover, the pairwise correlation analysis showed that, in fact, ORF1ab:A260V (NSP2), ORF1ab:S1188L (NSP3), ORF1ab:T1322K (NSP3), ORF1ab:K1795Q (NSP3), A28272G, ORF1ab:H1213Y (NSP13), N:P383L and ORF3a:Q213K had pairwise correlations above 0.85 (Figure 10). In addition, these variants formed a cluster that also included ORF8:I121L and ORF1ab:P970S (NSP13) (Figure 10B).

**Figure 10.**
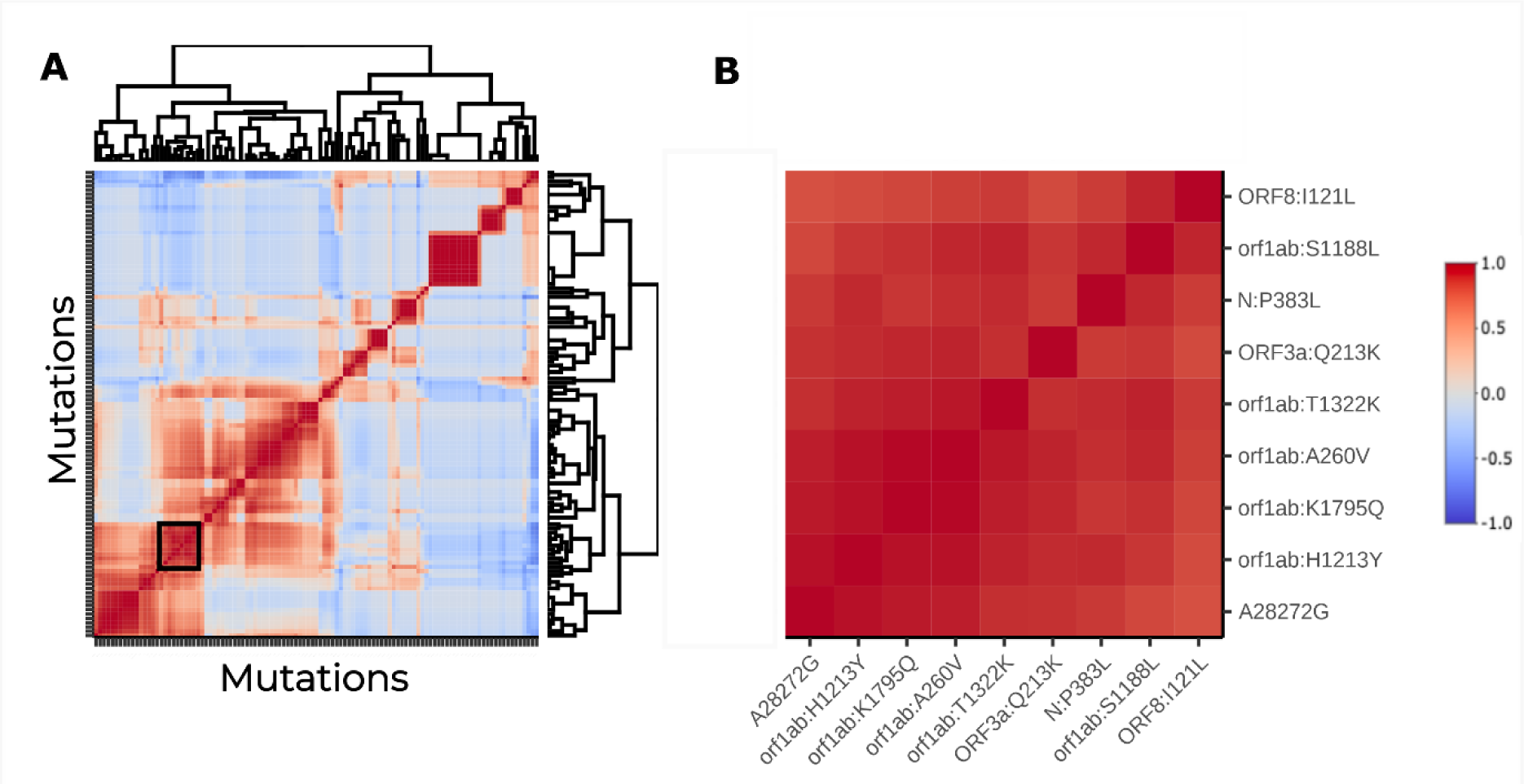
**Analysis of the association between polymorphism trajectories in the case study**. A) Hierarchically clustered heatmap of the pairwise Pearson’s correlation coefficients between the time series of allele frequencies in the case study. The cluster containing the previously found mutations is squared in black. B) Subset of the correlation heatmap, restricted to the cluster marked in (A).

#### Selective pressure

We calculated the number of non-synonymous substitutions per non-synonymous site (dN) and the number of synonymous substitutions per synonymous site (dS) for each sample. Despite of dN and dS being 0 in the first sample, dN showed a higher growth over time reaching a value of around 0.0007 in the last sample while dS kept a lower value of 0.0001, hinting at positive selection during the infection. The dN/dS ratio (ω) ranged between 1.11 and 5.98, with an average value of 2.36 (Figure 7B). These findings suggested a sustained positive selective pressure throughout the infection.

## Discussion

Chronic infections are becoming an important issue in SARS-CoV-2 evolutionary studies due to the relationship between the prolonged within-host viral evolution and the emergence of VOCs [20]. However, the study of serially-sampled SARS-CoV-2 samples lacks integrated workflows that facilitate the analyses. To close this gap, we have developed VIPERA, a tool that automatizes the analysis of serially-sampled SARS-CoV-2 samples.

A key strength of VIPERA is the combined use of phylogenetic and population genomics approaches to analyze SARS-CoV-2 samples and yield information to ascertain whether there is a serially-sampled infection or not. To do so, mapped reads are used in different ways to take into account the entire intra-host viral population. First, the lineage assignment of the samples is calculated using allele frequencies.

This analysis enables the user to detect co-infections or viral lineage replacement events, which can go unnoticed in a consensus genome analysis. Second, VIPERA also reports a maximum-likelihood phylogeny including the study and the context dataset. The tree allows the user to assess whether the studied samples are monophyletic, which is a good indicator for serially-sampled infections. Third, because nucleotide diversity is expected to be reduced for SARS-CoV-2 sequences from the same infection compared to independent samples, we use this metric to evaluate serially sampled infections. Comparison of within and between-host diversity has been previously used for viral outbreak analysis to detect transmission chains [21], and it has proven to be a strong indicator of serially-sampled infection in this work. Even when the context dataset includes some samples from the same patient as the studied sequences, we found that nucleotide diversity still contains enough signal to differentiate intra-patient variation. This is partly due to the robustness of the context dataset. Although VIPERA cannot assess in a systematic manner whether all samples in the context dataset are independent, we found identical results when we compared a customized context dataset with truly independent sequences and the automatic one. Thus, these results support the robustness of our approach to select a context dataset automatically. Finally, a strong temporal signal can further indicate that a target dataset has been serially sampled from a single infection, but it is not sufficient. Samples from different origins can exhibit a similar rate of evolution if they share collection dates, sampling locations and viral lineage. That could explain why our negative control showed a strong temporal signal. Furthermore, the size disparity between the two datasets in our negative control could influence too, because the larger dataset might be overshadowing the temporal signal of the smaller one. For this reason, temporal signals by themselves cannot be considered as evidence of intra-host evolution and must be taken into account only when previous evidence suggests a serially-sampled infection.

Once assessed if all sequences derive from the same infection, VIPERA’s results can be used to study the evolutionary process. Phylodynamic processes of inter-host and intra-host evolutionary dynamics can produce distinctive phylogenetic patterns [22]. In our work, monitoring the evolution of the virus during eight months allowed for the observation of both intra and between-host phylodynamic patterns within the same phylogeny. We achieved this by including a well-designed context dataset, as described earlier. We observed a balanced phylogeny for population level samples of our case study, but a heavily unbalanced one for within-patient samples, reflecting the different intra-host versus inter-host processes. VIPERA also reports dN/dS estimates through time which can reveal if natural selection has operated on the viral genomes during the studied serially-sampled infection. In the case studied here, dN/dS increased over time, showing a maximum value after eight months of infection. The phylogeny patterns along with the analysis of strength and mode of natural selection, suggests that intra-host evolution in our case study is driven by strong positive selection, and supports the hypothesis of a high evolutionary rate at the within-patient level.

Description of the intra-host nucleotide variants and their relationship with other variables such as collection date or other intra-host nucleotide variants is also reported by VIPERA. In our case study, we detected different mutations that are concerning because of their relationship with immune system evasion, such as ORF1ab:T1638I (NSP3), ORF1ab:S1188L (NSP3) and ORF3a:Q213K [23,24]. We also found mutations previously found in within-host evolution analyses such as N:P383L, ORF1ab:H1213Y (NSP13) and S:V143D + ΔY144 [25–27].

In summary, VIPERA facilitates the analysis of SARS-CoV-2 chronic infections by providing evidence for serially-sampled infection, describing the viral within-host evolution, and setting up an environment with the files needed for further custom within-host viral evolution analysis. For these reasons, we foresee VIPERA as an enhancer for SARS-CoV-2 serially-sampled infections studies and thus, helping to the surveillance of VOCs and to understand the mechanisms behind VOCs appearing. Although VIPERA is designed for reporting on SARS-CoV-2 sequence data, the framework could be extended to other viruses in further iterations of the software.

## Conclusions

VIPERA (Viral Intra-Patient Evolutionary Reporting and Analysis) is a new bioinformatic tool for studying and analyzing serially sampled SARS-CoV-2 infections. VIPERA provides an aggregate of analysis for detecting whether there is a serially-sampled infection or not, including novel approaches such as genetic diversity and genetic distance at the population level approaches. It also provides a description of the within-host evolution observed in the studied samples. Having undergone rigorous validation through two stringent control cases, our tool has proven its efficacy in a real-world case study. Being on the cusp of a new era in understanding the intra-host evolution of SARS-CoV-2, VIPERA paves the way for a more efficient analysis of serially-sampled SARS-CoV-2 samples.

## Methods

### Pipeline implementation

To facilitate the study of SARS-CoV-2 within-host evolution using data from single-virus serially-sampled infections, we have implemented VIPERA (Viral Intra-Patient Evolutionary Reporting and Analysis), a user-friendly, customizable and reproducible workflow using Snakemake [28], R v4.1.3 [29] and Python v3.10 [30] in addition to other software listed in Additional file 1: Table S2. VIPERA enables the automated analysis of an arbitrary number of samples collected from a single patient at different time points after infection. VIPERA takes as input sorted BAM files, consensus sequences in FASTA format and also a metadata file with collection dates, locations and GISAID IDs. While our tool is suited for the computational capabilities of an average laptop, we leveraged Snakemake profiles to ensure seamless deployment in a high-performance computing (HPC) environment. On our cluster, we achieve a consistent run time of under 15 minutes, using one Intel(R) Xeon(R) Gold 6230 CPU @ 2.10GHz and less than 1 GB of RAM. The run time decreases by up to a factor of 5 on 16 cores, using around 6 GB of RAM. The main output of VIPERA is a report file in HTML format that includes different analytical results and data visualization for detecting single-virus sustained infections and studying within-host evolution.

### Dataset retrieval and preprocessing

Three sets of SARS-CoV-2 samples were used in order to test and use VIPERA: a positive control, a negative control and a novel case.

For the positive control, we used 30 SARS-CoV-2 samples collected in Connecticut between June 1, 2021, and March 7, 2022, described as a chronic infection by Chrispin Chaguza et al. [11]. FASTQ files were fetched from the SRA using *fastq-dump*, implemented in the SRA toolkit v3.0.0 [31]. Reads were mapped against the Wuhan-Hu-1 reference genome (NCBI RefSeq accession no. NC_045512.2) [32] using BWA-MEM v0.7.17 [33]. ARTIC v4.1 primer schemes [34] were trimmed from the generated BAM files using *iVar* v1.4.2 [35]. Using *samtools* v1.17 [36] and *iVar* v1.4.2 [35] trimmed BAM files were sorted and indexed to obtain the consensus sequence with a minimum frequency threshold of 0.6 and a minimum depth of 20 reads.

The negative control and the novel case datasets were selected from samples for which we had access to BAM files, consensus sequences and metadata via the SeqCOVID Consortium. Viral samples were collected in the Hospital Clínic de Barcelona and sequenced in the Institute of Biomedicine of Valencia using the ARTIC v3 primer scheme [34]. Libraries were prepared using the Nextera Flex DNA Library Preparation Kit and sequenced on the Illumina MiSeq platform. Reads were processed through the SeqCOVID pipeline for SARS-CoV-2 bioinformatic analysis [37]. The case study comprised 12 samples collected from the same patient (Patient A) in Barcelona, Spain between March 30, 2020, and November 11, 2020, and previously designated as lineage B.1 (Table 1). For the negative control, the previous 12 samples were mixed with three samples from a different patient (Patient B), also collected in Barcelona, Spain between March 30, 2020, and November 16, 2020, and previously designated as (Table 1).

**Table 1.** Summary of the SARS-CoV-2 genomes analyzed in the negative control and in the case study. . The “NC” and “CS” abbreviations refer to the negative control and case study datasets, respectively. Index columns refer to the temporal order within each dataset, used as a label in neighbor-joining trees. Patient A is the target of our novel case study.

| ENA accession number | Collection date | Patient | NC ID | NC Index | CS ID | CS Index |
| --- | --- | --- | --- | --- | --- | --- |
| ERR5709045 | 2020-03-24 | A | NC_1 | 1 | CS_1 | 1 |
| ERR5709316 | 2020-04-01 | B | NC_2 | 2 | - | - |
| ERR5708640 | 2020-04-28 | A | NC_3 | 3 | CS_2 | 2 |
| ERR5709318 | 2020-05-18 | A | NC_4 | 4 | CS_3 | 3 |
| ERR5709345 | 2020-06-02 | A | NC_5 | 5 | CS_4 | 4 |
| ERR5709354 | 2020-06-22 | A | NC_6 | 6 | CS_5 | 5 |
| ERR5709412 | 2020-07-02 | B | NC_7 | 7 | - | - |
| ERR5709379 | 2020-08-03 | A | NC_8 | 8 | CS_6 | 6 |
| ERR5709385 | 2020-08-07 | A | NC_9 | 9 | CS_7 | 7 |
| ERR5709420 | 2020-08-19 | A | NC_10 | 10 | CS_8 | 8 |
| ERR5709429 | 2020-08-28 | B | NC_11 | 11 | - | - |
| ERR5708628 | 2020-11-06 | A | NC_12 | 12 | CS_9 | 9 |
| ERR5708657 | 2020-11-10 | A | NC_13 | 13 | CS_10 | 10 |
| ERR5709055 | 2020-11-12 | A | NC_14 | 14 | CS_11 | 11 |
| ERR5709463 | 2020-11-16 | A | NC_15 | 15 | CS_12 | 12 |

### Characterizing serially-sampled infections from a single virus

#### Longitudinal analysis of viral lineage assignment and admixture

The descriptive analysis of the target dataset of intra-patient samples includes the assignment of a Pango lineage according to sample consensus sequences, as well as the evaluation of possible lineage admixture within each sample. A lineage is assigned to the genome sequences of each sample using Pangolin v4.3 [38] in accurate (UShER) mode. A demixing step is performed using Freyja v1.4.2 [39], which utilizes read mappings to estimate the lineage admixture of each sample based on lineage-defining mutational barcodes by solving a convex optimization problem.

#### Construction of a context dataset

The analyses require a collection of independent samples —ideally, samples that originate from different hosts and separate infection events. This set of samples is referred to as the “context dataset” in our study. Automated construction of the context dataset is enabled by default, contingent upon the provision of user credentials for the GISAID SARS-CoV-2 database [40], using *GISAIDR* v0.9.9 [41]. This facilitates the retrieval of a dataset comprising samples that fulfill the spatial, temporal and phylogenetic criteria, including a sampling location that corresponds to that of the target samples, a collection date that falls within a time window encompassing 95% of the date distribution of the target samples (with 2.5% trimmed at each end to account for extreme values) ± 2 weeks, and a lineage assignment that is shared by at least one of the target samples. During the process, a series of tweakable checkpoints are enforced to ensure a robust downstream analysis. By default, samples whose GISAID accession number matches any of the target samples are removed. In addition, the dataset is rejected if the number of samples does not allow at least as many possible combinations as replicates. Alternatively, a manually constructed context dataset may be provided. For all the analyses shown in this article, an automatically constructed context dataset has been used. Additionally, a manually constructed context dataset was also used for the case study to compare the results with the ones obtained using an automatically constructed context dataset.

#### Nucleotide diversity comparison

Nucleotide diversity (π) of the target dataset is compared with that of the context dataset, composed of independent samples. By default, nucleotide diversity is calculated for 1000 random sample subsets of size equal to the number of target samples, extracted with replacement from the context dataset. The number of replicates can be easily modified by the user. Then, the obtained distribution is compared with the nucleotide diversity obtained for the target dataset; empirically, if the π distribution is not normal, or via parametric tests, if it is. Calculations are performed in R, and nucleotide diversities are calculated with *pegas* v1.2 [42].

#### Assessing phylogenetic relationships and temporal signal

Consensus sequences of the target and context datasets are aligned to the Wuhan-Hu-1 reference genome (NCBI RefSeq accession number: NC_045512.2) [32] using Nextalign v2.13 [14]. Positions classified as problematic [43] are masked in the alignments. Then, a maximum-likelihood phylogeny is constructed using IQTREE v2.2.2.3 [44]. By default, inference is performed under a GTR substitution model with empirical base frequencies, a heterogeneity model with a proportion of invariable sites and a discrete Gamma distribution with 4 rate categories, ultrafast bootstrap (UFBoot) [45,46] with 1000 replicates, and the Shimodaira–Hasegawa-like approximate likelihood ratio test (SH-aLRT) [47] with 1000 replicates. This inference enables the study of the taxonomic grouping of the target dataset within the relevant epidemic context.

To take the within-host variability in the viral population into account, we propose a pairwise distance metric between samples that integrates the differences in allele frequencies across the whole genome.

We define the difference between two vectors of *J* allele frequencies, based on the FST measure [48], such that the distance between two samples (*M* and *N*) is the sum for all *I* polymorphic sites of the differences between allele frequencies at each position (see Equation 1). Then, with this distance matrix, a neighbor-joining tree is constructed in R using *ape* v5.7 [49]. Patristic distances to the root are calculated with *adephylo* v1.1-13 [50].

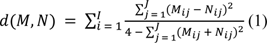

Finally, the evolutionary rate is estimated by linear regression of the patristic distances to the root in each phylogeny on the days passed since the first within-patient sample collection, using the *lm* implementation in the *stats* R library.

### Describing within-host variability

#### Variant calling and nucleotide variant description

Variants are called using *samtools* v1.17 [36] and *iVar* v1.4.2 [35] using a reconstructed ancestral genome as reference to restrict the analysis to sequence variation related to the within-host evolution. Variants are re-annotated using *snpEff* v5.1d [51]. To reconstruct the ancestral sequence, the target samples are aligned to the Wuhan-Hu-1 reference genome (NCBI RefSeq accession no. NC_045512.2) [32] using Nextalign v2.13 [14]. Then, the ancestral genome is obtained with IQTREE v2.2.2.3 [44]. By default, maximum-likelihood trees are inferred under a GTR substitution model with empirical base frequencies and a heterogeneity model with a proportion of invariable sites and a discrete Gamma distribution with 4 rate categories. The quality criteria for variant calling were a minimum base quality of 20, a minimum depth of 30 and a minimum frequency cutoff of 5%. Nucleotide variants supported by less than 20 reads or less than 2 reads in one strand were filtered out.

The distribution for the polymorphisms found along the SARS-CoV-2 genome is calculated using a sliding window (default width: 1000 nucleotides; step: 50 nucleotides). The number of mutations per site for each window is represented on its right side. Positions are annotated using the Python library *gb2seq* v0.0.20 [52].

To select the most interesting polymorphisms to plot, we perform a linear regression of the allele frequencies of each polymorphism on the time (in days) elapsed since the first within-patient sample collection. Correlation is measured with the Pearson’s correlation coefficient, and the p-value of the linear regression is adjusted for multiple testing using the Benjamini-Hochberg method [53]. This analysis is performed using the *stats* R library. Then, polymorphisms that have a significant correlation with time progression are selected for further characterization. Additionally, sites with more than one alternative allele are also selected to monitor potential associations or interactions between the alternative alleles.

Moreover, we calculate pairwise correlations between allele frequencies for all pairs of polymorphisms. Mutations are hierarchically clustered based on correlation values. Pairwise correlations are measured with the Pearson’s correlation coefficient using the *stats* R library. Display of the hierarchical clustering and correlation values is carried out through the *heatmaply* R library [54] with *hclust* (from the *stats* R library) as the clustering function.

#### Investigating traces of selection

To track selection footprints, substitutions per synonymous site (dS) and substitutions per non-synonymous site (dN) are calculated for each sample. Synonymous and non-synonymous sites are calculated with respect to the reconstructed ancestral sequence. Then, dN and dS are calculated taking into account allele frequencies. Calculations are performed in Python using the Nei-Gojobori method [55] with support of *gb2seq* v0.0.20 [52] for codon annotation.

## Supporting information

Additional File 2: variant calling results

Additional File 3: VIPERA report

## Declarations

### Availability of data and materials

VIPERA is a cross-platform Snakemake (≥7.19) workflow written in Python and R, released as open-source software under the GNU GPLv3 license. Source code is available in GitHub (https://github.com/PathoGenOmics-Lab/VIPERA, release v1.0.0). The VIPERA report of the case study dataset is available as Additional File 3.

Sequencing data from the positive control is available through its source publication by Chaguza et al. [11]. Raw sequencing data from the negative control and the novel case study are available at the ENA. Accession numbers are provided in Table 1. Read mappings and consensus genomes can be accessed via DOI: 10.20350/digitalCSIC/15648.

### Competing interests

The authors declare that they have no competing interests.

### Funding

This work was funded by the Spanish Ministry of Science (CNS2022-135116) and the European Commission – NextGenerationEU (Regulation EU 2020/2094), through the Global Health Platform (PTI+ Salud Global) of the Spanish National Research Council (CSIC). MAH and PRR are supported by the PTI+ Salud Global. JS is supported by the CSIC’s JAE intro programme. FGC was funded by project PID2021-127010OB-I00 from the Spanish Ministry of Science and CIPROM2021-053 from the Generalitat Valenciana. IC is funded by the European Research Council (101001038-TB-RECONNECT) and the Spanish Ministry of Economy, Industry and Competitiveness (PID2019- 104477RB-I00). MC is funded by the Spanish Ministry of Science (PID2021-123443OB-I00).

### Authors’ contributions

JS: conceptualization, data curation, investigation, methodology, software, formal analysis, validation, visualization, writing – original draft. MAH: conceptualization, data curation, investigation, methodology, software, formal analysis, validation, writing – original draft. PRR: software, writing – review & editing. AV, JV, PCJ, IC: resources. FGC: funding acquisition, writing - review & editing. MC: conceptualization, methodology, project management, writing - review & editing, funding acquisition, supervision.

#### Acknowledgements

The computations were performed on the HPC cluster Garnatxa, at the Institute for Integrative Systems Biology (I^2^SysBio), a joint collaborative research institute involving the University of Valencia (UV) and the Spanish National Research Council (CSIC). We thank Dr. Anne Hahn and Dr. Nathan Grubaugh (Laboratory of Epidemiology of Public Health, Yale School of Public Health, USA) for sharing the information about the sequences we identified as belonging to a previous study. We also acknowledge the SeqCOVID Consortium for providing the sequencing data of the negative control dataset and the case study.

## Notes

https://doi.org/10.20350/digitalCSIC/15648

https://github.com/PathoGenOmics-Lab/VIPERA

### Competing Interest Statement

The authors have declared no competing interest.

