## Additional File 1: supplemental tables for "VIPERA: Viral Intra-Patient Evolution Reporting and Analysis"

**Table S1. Intermediate files produced by VIPERA.** File paths are relative to the output directory, which is defined via the VIPERA configuration files. Names between brackets are defined via these files as well.

| FILE PATH | FILE DESCRIPTION |
|---|---|
| nextalign/{output name}.masked.filtered.fasta | Masked alignment of the target dataset |
| {output name}.lineage_report.csv | Dataset lineage report, calculated with Pangolin |
| tree/{output name}.treefile | Maximum-likelihood phylogeny of the target dataset in Newick format |
| demixing/{sample name}/{sample name}.variants.tsv | Variant calling results of each sample against the reference genome |
| demixing/{sample name}/{sample name}.demixed.tsv | Raw demixing results of each sample |
| {output name}.freyja_demixing.csv | Dataset lineage demixing summary |
| {output name}.ancestor.fasta | Maximum-likelihood ancestral sequence reconstruction of the MRCA for the target dataset in FASTA format |
| {output name}.masked.filtered.tsv | Variant calling results against the dataset ancestor |
| context/nextalign/context_sequences.aligned.masked.fasta | Masked alignment of the context dataset |
| context/duplicate_accession_ids.txt | List of GISAID IDs removed from the context dataset due to their accession ID being already in the target dataset |
| tree_context/{output name}.treefile | Maximum-likelihood phylogeny of the target dataset within its spatiotemporal context in Newick format |
| report/tables/*.csv | Data for creating the report figures |
| report/plots/*.png | Figures generated for the report file |

**Table S2. VIPERA dependencies.** These include programming languages (in italics), computer programmes (in bold) and code libraries.

| SOFTWARE | LIBRARY | VERSION | REFERENCE |
|---|---|---|---|
| **Snakemake** | | **≥ 7.19** | 10.12688/f1000research.29032.2 |
| *R* | | *≥ 4.1.3* | www.R-project.org |
| *R* | tidyverse | 2.0.0 | 10.21105/joss.01686 |
| *R* | ggrepel | 0.9.3 | CRAN.R-project.org/package=ggrepel |
| *R* | quarto | 1.2 | CRAN.R-project.org/package=quarto |
| *R* | stringi | 1.7.12 | 10.18637/jss.v103.i02 |
| *R* | ggpubr | 0.6.0 | CRAN.R-project.org/package=ggpubr |
| *R* | ggtree | 3.2.0 | 10.1201/9781003279242 |
| *R* | ape | 5.7 | 10.1093/bioinformatics/bty633 |
| *R* | adephylo | 1.1_13 | 10.1093/bioinformatics/btq292 |
| *R* | pegas | 1.2 | 10.1093/bioinformatics/btp696 |
| *R* | data.table | 1.14.8 | CRAN.R-project.org/package=data.table |
| *R* | future.apply | 1.11.0 | 10.32614/RJ-2021-048 |
| *R* | scales | 1.2.1 | CRAN.R-project.org/package=scales |
| *R* | showtext | 0.9_6 | CRAN.R-project.org/package=showtext |
| *R* | jsonlite | 1.8.5 | arxiv.org/abs/1403.2805 |
| *R* | logger | 0.2.2 | CRAN.R-project.org/package=logger |
| *R* | gt | 0.9.0 | CRAN.R-project.org/package=gt |
| *R* | heatmaply | 1.4.2 | 10.1093/bioinformatics/btx657 |
| *R* | readr | 2.1.4 | CRAN.R-project.org/package=readr |
| *R* | devtools | 2.4.5 | CRAN.R-project.org/package=devtools |
| *R* | GISAIDR | 0.9.9 | 10.5281/zenodo.6474693 |
| *Python* | | *3.10* | https://www.python.org/ |
| *Python* | biopython | 1.81 | 10.1093/bioinformatics/btp163 |
| *Python* | pandas | 2.0.3 | 10.5281/zenodo.3509134 |
| *Python* | pip | 23.2.1 | https://pypi.org |
| *Python* | gb2seq | 0.2.20 | github.com/virologycharite/gb2seq |
| **mafft** | | **7.520** | 10.1093/molbev/mst010 |
| **entrez-direct** | | **16.2** | ncbi.nlm.nih.gov/books/NBK179288 |
| **freyja** | | **1.4.2** | github.com/andersen-lab/Freyja |
| **iqtree** | | **2.2.2.3** | 10.1093/molbev/msaa015 |
| **nextalign** | | **2.13** | 10.1093/bioinformatics/bty407 |
| **pangolin** | | **4.3** | 10.1093/ve/veab064 |
| **quarto** | | **1.3.450** | github.com/quarto-dev/quarto-cli |
| **iVar** | | **1.4.2** | 10.1186/s13059-018-1618-7 |
| **samtools** | | **1.17** | 10.1093/gigascience/giab008 |
| **snpEff** | | **5.1d** | 10.4161/fly.19695 |
