## Additional File 3: VIPERA report for "VIPERA: Viral Intra-Patient Evolution Reporting and Analysis": additional_file_3.html

VIPERA report: case\_study


### VIPERA report: case\_study

Workflow version: v1.0.0

Published

2023-10-23

#### Summary

- 1 Summary of the target samples dataset
- 2 Evidence for single, serially-sampled infection
  - 2.1 Lineage admixture
  - 2.2 Phylogeny and temporal signal
  - 2.3 Nucleotide diversity comparison
- 3 Evolutionary trajectory of the serially-sampled SARS-CoV-2 infection
  - 3.1 Number of polymorphic sites
  - 3.2 Description for intra-host nucleotide variants
  - 3.3 Time dependency for the intra-host nucleotide variants
  - 3.4 Correlation between alternative alleles
  - 3.5 Non-synonymous to synonymous rate ratio over time

 

#### 1 Summary of the target samples dataset

Table 1:

Target dataset summary

| Sample | Index | Collection Date | Lineage |
| --- | --- | --- | --- |
| ERR5709045 | 1 | 2020-03-24 | B.1 |
| ERR5708640 | 2 | 2020-04-28 | B.1 |
| ERR5709318 | 3 | 2020-05-18 | B.1 |
| ERR5709345 | 4 | 2020-06-02 | B.1 |
| ERR5709354 | 5 | 2020-06-22 | B.1 |
| ERR5709379 | 6 | 2020-08-03 | B.1 |
| ERR5709385 | 7 | 2020-08-07 | B.1 |
| ERR5709420 | 8 | 2020-08-19 | B.1 |
| ERR5708628 | 9 | 2020-11-06 | B.1 |
| ERR5708657 | 10 | 2020-11-10 | B.1 |
| ERR5709055 | 11 | 2020-11-12 | B.1 |
| ERR5709463 | 12 | 2020-11-16 | B.1 |

#### 2 Evidence for single, serially-sampled infection

##### 2.1 Lineage admixture

The estimated lineage admixture for each sample has been calculated using Freyja.

Figure 1: Estimated lineage admixture of each sample. Samples in the X-axis are ordered chronologically, from more ancient to newer.

##### 2.2 Phylogeny and temporal signal

A maximum likelihood tree of the target and context samples has been build using IQTREE. The target samples are monophyletic. The clade that contains all the target samples is supported by a **UFBoot** score of \(92\)% and a **SH-aLRT** score of \(92.7\)% (Figure 2).

Figure 2: Maximum-likelihood phylogeny with 1000 support replicates of target datasets and their context samples. The clade that contains the target samples is squared in red.

A neighbor-joining tree has been constructed using pairwise distances between target samples (Figure 3), based on the allele frequencies measured from read mappings.

Figure 3: Neighbor-joining tree based on the pairwise allele frequency-weighted distances.

Root-to-tip distances measured on this tree have been correlated with time, obtaining a \(R^2\) of **0.9459** and a p-value of \(< 0.001\). The estimated substitution rate is **32.02** substitutions per year (Figure 4).

Figure 4: Scatterplot depicting the relationship between root-to-tip distances and the number of days passed since the first sample. The red line shows the linear model fit.

##### 2.3 Nucleotide diversity comparison

Nucleotide diversity (π) has been calculated for \(1000\) random sample subsets of size \(12\), extracted with replacement from the context dataset. The distribution of the nuclotide diversity is assumed to not be normal after performing a Shapiro-Wilk test (p-value of \(< 0.001\)).

The nucleotide diversity of the target samples is \(4.111673e-05\) (red line in Figure 5). Assuming the independence of the context samples, the empirical p-value of the context samples having a nucleotide diversity (in orange in Figure 5) as low as that of the target dataset is \(< 0.001\).

Figure 5: Analysis of the nucleotide diversity (π). The orange line describes a normal distribution with the same mean and standard deviation as the distribution of π from \(1000\) subsets of \(12\) sequences from the context. The red vertical line indicates the π value of the target samples.

#### 3 Evolutionary trajectory of the serially-sampled SARS-CoV-2 infection

##### 3.1 Number of polymorphic sites

Sites with minor allele frequency > 0.05 are considered polymorphic. The linear association between the collection date of the samples and the number of polymorphic sites has an \(R^2\) of **0.7172** and a p-value of \(< 0.001\) (Figure 6).

Figure 6: Number of polymorphic sites along time. The blue line shows the linear model fit.

##### 3.2 Description for intra-host nucleotide variants

A total of 99 different single nucleotide variants (SNV) and 10 insertions and deletions (indels) have been detected along the genome (Figure 7).

- Whole genome
- Spike ORF

Figure 7: Summary of the intra-host accumulation of nucleotide variants, using the reconstructed dataset ancestor as reference. A) Nucleotide variants per site along the SARS-CoV-2 genome. Relative abundance of NVs is calculated with a sliding window of width \(1000\) nucleotides and a step of \(50\). Labels indicate the coding regions of the non structural proteins (NSP) within ORF1ab. B) Genome variation along the genome for each sample. The Y-axis displays samples in chronological order, with the earliest collection date at the bottom, and the latest, at the top.

Figure 8: Summary of the intra-host accumulation of nucleotide variants in the spike sequence, using the reconstructed dataset ancestor as reference. A) Nucleotide variants per site along the S gene. Relative abundance of NVs is calculated with a sliding window of width \(1000\) nucleotides and a step of \(50\). B) Genome variation along the S gene for each sample. The Y-axis displays samples in chronological order, with the earliest collection date at the bottom, and the latest, at the top.

##### 3.3 Time dependency for the intra-host nucleotide variants

The correlation of the allele frequency of each NV with the time since the initial sampling has been calculated (Figure 9).

Figure 9: Pearson’s correlation coefficients and adjusted p-values of allele frequencies with time. Red dashed line indicates adjusted \(p = 0.05\). Labeled dots represent nucleotide variants correlated with time (adjusted \(p < 0.05\)).

Significantly correlated nucleotide variants are described in more detail in Figure 10.

Figure 10: Time series of relative allele frequencies. The shown positions include nucleotide variants with a significant correlation with time and sites with more than two possible states. Each subplot depicts the progression of the allele frequencies in time for a given genome position.

##### 3.4 Correlation between alternative alleles

To detect possible interactions between mutations, pairwise correlation between allele frequencies are calculated (Figure 11). The heatmap is an interactive figure that allows zooming in on specific regions.

Figure 11: Interactive hierarchically clustered heatmap of the pairwise Pearson’s correlation coefficients between the time series of allele frequencies in the case study.

##### 3.5 Non-synonymous to synonymous rate ratio over time

To track selection footprints, the substitutions per synonymous site (\(dS\)) and per non-synonymous site (\(dN\)) for each sample with respect to the reconstructed ancestral sequence have been calculated (Figure 12), as well as their ratio (\(\omega = dN/dS\); Figure 13).

- \(dN\) and \(dS\)
- \(\omega\) (\(dN/dS\))

Figure 12: Time series of dN and dS. Each point corresponds to a different sample, sorted in chronological order.

Figure 13: Time series of \(\omega\) (\(dN/dS\)). Each point corresponds to a different sample, sorted in chronological order.
